# Massively parallel quantification of phenotypic heterogeneity in single cell drug responses

**DOI:** 10.1101/2020.12.18.423559

**Authors:** Benjamin B. Yellen, Jon S. Zawistowski, Eric A. Czech, Caleb I. Sanford, Elliott D. SoRelle, Micah A. Luftig, Zachary G. Forbes, Kris C. Wood, Jeff Hammerbacher

## Abstract

Single cell analysis tools have made significant advances in characterizing genomic heterogeneity, however tools for measuring phenotypic heterogeneity have lagged due to the increased difficulty of handling live biology. Here, we report a single cell phenotyping tool capable of measuring image-based clonal properties at scales approaching 100,000 clones per experiment. These advances are achieved by exploiting a novel flow regime in ladder microfluidic networks that, under appropriate conditions, yield a mathematically perfect cell trap. Machine learning and computer vision tools are used to control the imaging hardware and analyze the cellular phenotypic parameters within these images. Using this platform, we quantified the responses of tens of thousands of single cell-derived acute myeloid leukemia (AML) clones to targeted therapy, identifying rare resistance and morphological phenotypes at frequencies down to 0.05%. This approach can be extended to higher-level cellular architectures such as cell pairs and organoids and on-chip live-cell fluorescence assays.

## Introduction

In recent years, advances in sequencing technology have enabled deep, clonally resolved views into the genomic and transcriptional heterogeneity that exists within cellular populations.[1-4] This variance is important, as it likely drives much of the phenotypic heterogeneity that underpins physiological and pathological programs.[3-5] While single cell genomic tools can now routinely measure the mutational or transcriptional profiles of >100,000 individual cells in a single experiment,[6,7] similar tools for measuring single cell phenotypic heterogeneity and dynamics remain elusive owing to the complexities of working with live biology. One promising approach for capturing phenotypic heterogeneity on a massive scale entails organizing a high-density array of individual cells that can be continuously observed over time microscopically, with or without chemical or physical perturbations. Imaging these isolated clones can reveal phenotypic distributions, including rare phenotypes of biological significance, such as cells that respond uniquely to important stimuli or produce distinct secreted factors.[8,9] However, to date no existing platforms have demonstrated the ability to measure single cell phenotypes at throughputs approaching the 100,000 clone scale. The only platform that approaches this benchmark is the Berkeley Lights Beacon® instrument, but to our knowledge that platform is currently unable to perform more than four parallel experiments per instrument, limiting the ability to analyze phenotypic responses of diverse cell types to assorted stimuli.[9]

In cancer, rare clones that survive in the presence of chemotherapeutic treatments often drive recurrence of drug resistant disease.[10,11] In the laboratory, these clones have traditionally been isolated and studied individually through a weeks- to months-long process of selection, enrichment, and clonal isolation. As a result, it has been difficult to quantify the abundance of resistant clones in a population, directly define clonal growth properties, or scale analyses to different tumor samples, cell lines, drugs, and doses. Given the complexity and heterogeneity of resistant clones within individual patients, it is expected that new, integrative approaches will be necessary to design drug therapies capable of suppressing the collective growth of resistant subclonal populations. This necessity underscores the importance of technologies that can measure these properties at scale.[10-12]

In this report, we present the first single cell phenotyping platform that can reach the scale of 100,000 clones in a single, multi-day, time-resolved experiment, all performed in parallel by one instrument. Our approach is made possible by fundamental advances in microfluidic chip design, improvements in methods for long-term culturing and microscopic observation of single cells, and finally advances in image processing and analysis software that allow for large image-based datasets to be automatically analyzed down to the level of individual cell morphology. Specifically, we report on a novel microfluidic design that represents the most efficient microfluidic trapping architecture to date, and we also demonstrate robust, cost-effective methods for maintaining mammalian cells on-chip over sufficient time to identify rare phenotypic properties like drug resistance. These results pave the way for more efficient methods for credentialing drugs and ultimately, improved selection of therapeutic regimens for patients. More broadly, by enabling flexible phenotypic single cell profiling at massive scale, this platform may facilitate the functional characterization of diverse and complex cellular populations.

## Results

### A novel microfluidic flow regime enables high efficiency single cell trapping

In our chip design, there are two unique flow regimes distinguished by the opposite directions of fluid flow that can exist in the rungs of a microfluidic ladder network, which is caused by the difference in fluidic resistances in the rails of the ladder (Fig. 1a, see supplementary theory for details). This two-state flow system is present in both microfluidic mesh and ladder networks; however prior works have all used only one flow regime in cell trapping devices [13-17] in which the resistance through the trap, R_A_, (one of the rails of the ladder) is higher than the hydrodynamic resistance through the bypass section, R_T_, (the opposite ladder rail). In previous studies, the fluid flow in the ladder rung points away from the trap (see the blue arrows in Fig. 1a,b), which leads to a less efficient trap since cells have the ability to bypass the traps by following the streamlines towards the serpentine section.

**Figure 1.**
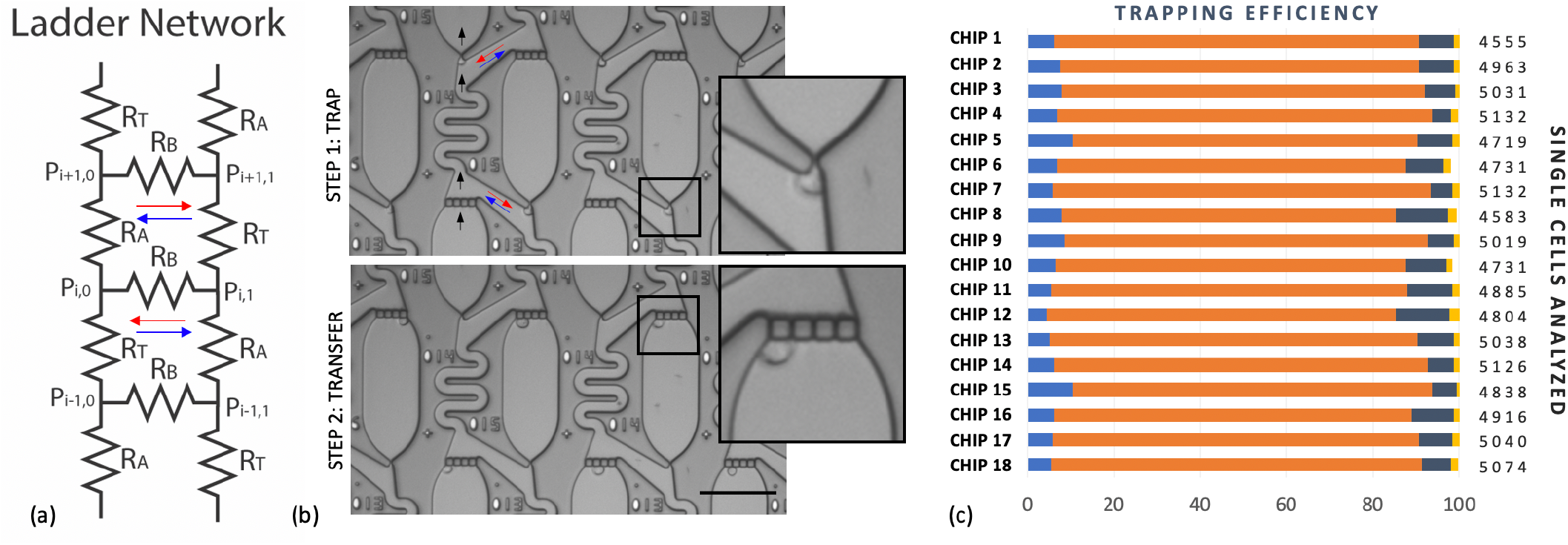
Working Principle. The microfluidic device can be modeled as a ladder resistor network (a), which has two flow regimes that have high (or low) cell trapping efficiency. The high (low) trapping flow regimes are depicted by the red (blue) arrows, respectively, in which the fluid joins (splits) before entering the apartment. The cells are first trapped in the constriction (b, top), and then transferred into the apartments with a brief pressure pulse (b, bottom). A higher magnification view of each step in the trap and transfer process is shown. The results from one experiment consisting of 18-chips (c) capturing a total of 88,317 cells shows that on average 83% of apartments contain a single cell (orange bars), 7% are empty (light blue bars), and 10% have more than one cell (dark blue bars [2 cells] and gold bars [> 2 cells]). Scale bar is 100-µm.

Surprisingly, there have been no prior reports of microfluidic devices in the other flow regime, R_A_ < R_T_, which is vastly more efficient as a cell trap because, in this case, the flow joins rather than splits at the entrance to the trap (see the red arrows in Fig. 1a,b). This improved trapping efficiency comes at the expense of a larger device footprint, since it is necessary to design longer and narrower channels for the bypass section in order to achieve the required resistance ratio. However, this approach is advantageous when the goal is to improve the trap occupancy rate and make more efficient use of limited cell samples.

The working method of this trapping approach is based on a self-limiting principle, in which the fluid flow is modulated by the cell’s physical presence in a trap, which functions as a switch to alternate between the two flow states. When the trap is initially empty, it is in a high efficiency flow state for capturing cells where R_A_ < R_T_. After a trap has captured a cell, the flow profile changes because the cell’s physical presence modifies the resistance through the trap, thus switching the flow state to the low efficiency capture state where R_A_ > R_T_. The high resistance of the occupied traps causes subsequent cells to bypass the occupied traps and diverts them downstream toward unoccupied traps. As a result, the cells populate the array in a deterministic fashion with most of the traps becoming filled in the order that cells were introduced onto the chip.

To load the cells onto the chip, a cell suspension at a concentration of 10^6^ cells per mL is prepared in a 0.2 µm-filtered aliquot of cell culture media, and then a 10 – 20 µL aliquot of cell suspension is placed into the inlet reservoir, after which it takes approximately 3 to 5 minutes to fill all of the traps by applying negative pressure (20 to 50 mbar) at the microfluidic outlet. The remaining cells are then rinsed from the device by washing the inlet several times and then flowing clean filtered media through the chip for another minute, leading to a trapping distribution similar to that shown in Fig(1b, top).

Once the traps are filled, the cells are transferred into the apartments by applying a sub-second elevated pressure pulse, which squeezes the cells through the constrictions into the adjacent apartments. These mechanical perturbations are benign and have been successfully employed in various drug and gene delivery applications.[18-20]. As a general rule, we found that a 1:3 or 1:4 ratio for the width of the trap region compared to the diameter of the cell was ideal; this allowed cells to be consistently trapped and retained at low pressures (∼20 mbar), but reliably transferred into the apartments at higher pressures (∼500 mbar). In our chip designs, the single cell capture efficiency worked best when the front trap width is in the range 3 – 6 µm for cells for cell diameters in the range of 10 – 25 µm. Representative images of the cell positions during each step of the trap and transfer process are shown in Fig. 1b.

We also developed automated methods for reading the individual apartment addresses from the images and quantifying the number of cells in each chamber through brightfield image classification techniques. These classifiers are based on standard image segmentation models that have been trained to detect the instances of each cell in each chamber at each time point [21,22], as described in detail in the methods section. With this software package, we were able to quantify the trapping efficiency in the array and assess any spatial biases that were used to improve the microfluidic architecture.

In this platform, we are simultaneously optimizing two metrics of performance, namely (1) the number of traps that end up capturing a single cell (typically ∼80% in our hands), and (2) the number of cells needed to completely fill all of the traps, which is related to how the cells are distributed in each of the parallel channels during the loading process. Both of these parameters need to be optimized to effectively make use of limited cell samples. Since this design is very efficient at capturing cells, the 6,016 traps in the device are consistently filled when ∼10,000 cells are introduced to the inlet. Due to the combination of fabrication defects, presence of debris in the cell culture media, and incompletely dissociated cell suspensions, the trap occupancy rates for MOLM-13 cells was found on average to yield ∼80% single cells, ∼10% empty chambers, and ∼10% chambers having more than one cell (Fig. 1c).

### Cell pairs and reproducible cell clusters can be organized with high efficiency

This platform has the ability to organize other types of cellular architectures for myriad potential cellular analysis applications by tuning the device geometry and/or serially repeating the trap and transfer process. The ability to form heterogeneous cell pairs (Fig. 2a), for example, has potential applications in immunooncology and in forming different types of cellular micro-environments. A similar approach has been employed by others for fabricating hybridomas [13], and pairing T cells with other cells [23]. We demonstrate this pairing ability by first organizing an array of MOLM-13 cells that were labeled with CellTrace Far Red dye, and then repeating this process with the same cells that were instead loaded with CellTrace CFSE dye. In each step, we obtained ∼80% single cell capture efficiency, and this yielded an overall efficiency of ∼64% of chambers having heterogenous red/green single cell pairs.

**Figure 2.**
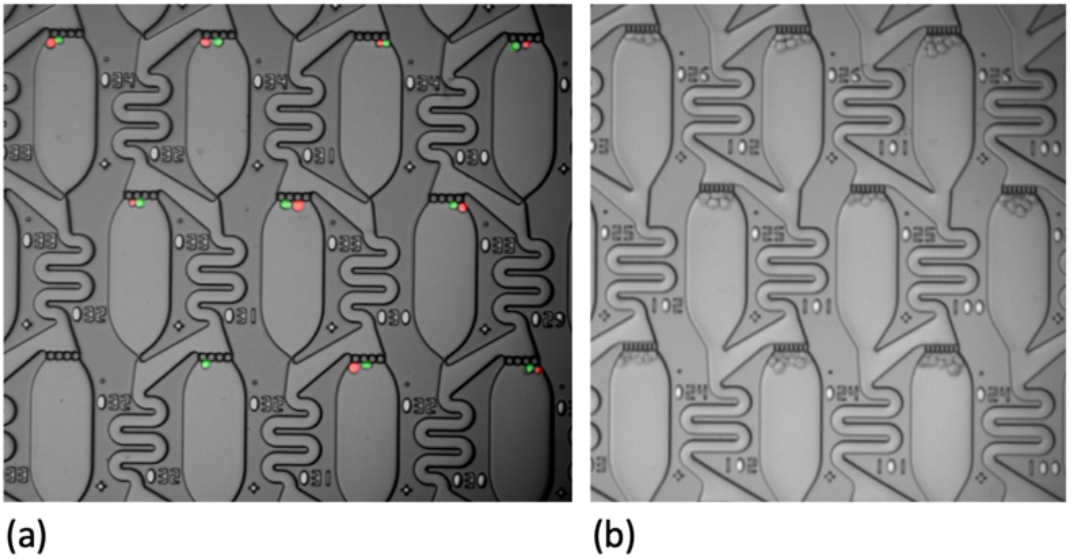
Multi-cellular architectures. (a) Heterogeneous single cell pairs are formed by repeating the trap and transfer process with different cell types, whereas (b) highly reproducible cell clusters are created in a single shot when the front trap is opened up.

This device can be modified to form reproducible cell groupings in a single shot by opening up the front trap, as shown in Fig. 2b. For this geometry, repeatable clusters of 6-10 cells per apartment were organized reliably across the entire chip, and this approach may have potential applications for rapidly creating spheroids or organoids in a highly parallel format.

### Rare cell phenotypes are observed in multi-day cell culture experiments

The chips are fabricated by deep reactive ion etching (DRIE) of Silicon wafers, then anodically bonding the wafers to glass lids, followed by dicing the wafers into individual chips, and finally assembling the chips into custom-machined microfluidic chip holder (see methods section for detailed fabrication process). This fabrication approach can be readily accomplished in a standard university cleanroom and allowed us to fabricate features as small as 2 µm. The microfluidic architecture was designed such that cells were able to squeeze through the front traps having 3 – 6 µm constrictions; however, cells were retained by the parallel frit structure at the back of the chambers that had smaller 2 µm constrictions. The geometry allowed fluid to pass through the apartments while retaining the cells inside the chambers over many days, enabling the study of clonal growth patterns and variability in morphological features.

To achieve steady perfusion of media into the chips while inside the incubator, we connect 10mL syringes to the inlet and outlet, which serve as media reservoirs. We fill the inlet syringe with media while the outlet syringe is connected to a vacuum line. In this way, we ensure that the media reservoir at the inlet has unimpeded gas exchange with the ambient conditions inside the incubator. We maintain good cell viability by using weak vacuum pressures in the range of -20 to -50 mbar to continuously flow media through the device. For this specific chip, an optimal flow rate of ∼5 mL/day is sufficient to remove metabolic waste products and provide fresh nutrients. However, the exact flow rates need to be tuned for other microfluidic geometries, cell types, and other experimental parameters.

The simplicity of this microfluidic design allows for re-sealable connections to be made easily between the chip and the external pressure controllers, which enables many on-chip experiments to be conducted in parallel (Fig. S1a). Each day the chip is disconnected from the pumping system to perform imaging on a standard fully automated microscope (Fig. S1b). To rapidly acquire high-resolution images of each apartment in the chip, we developed imaging algorithms that employ microscope image quality focus classifiers [24] and image-segmentation computer vision models (Mask-RCNN) [25] to identify fiducial marks on the chips and determine the optimal focus for each image (see methods section and github repository posted online [26]). As a compromise between image quality and speed, we opted to perform microscopy at 10X magnification, allowing the entire chip to be tiled with ∼300 images where 20 apartments are captured in each field of view. This approach allows us to image each chip within 5-10 minutes depending on the number of fluorescent channels, and it provided the bandwidth to image up to 18 chips per day (see Fig 1c and Fig S1).

With the ability to repeatedly image many chips in parallel over many days and analyze cell properties per chamber with our software pipeline, we have the statistical power to discover rare phenotypic variants of biological significance. For example, in Fig. 3a we plot the growth rate as a function of the time-averaged mean cell area across the 11,094 single cell clones that maintained positive growth rates over 96 hr. The distribution of growth rates in each of the three chips show similar phenotypic distributions, each displaying medians of ∼0.95 doublings/day with the middle quartiles falling in the range of 0.84 – 1.12 doublings/day and the fastest growth rate exceeding 1.5 doublings/day. Beyond growth measurements, we found an interesting subset of clonal cellular populations that not only were fast growers but also abnormally large compared to the bulk population. A few of these rare phenotypes were found in each chip, and their frequency in the parental line was assessed to be ∼0.05% for this cell line (See Fig. 3b). These rare cells consistently presented with a pear-shaped morphology (Fig. 3c), and the fact that they are larger across all timepoints and have similar morphologies provides intriguing evidence of (epi)genetically heritable cell size and shape regulation.[27]

**Figure 3.**
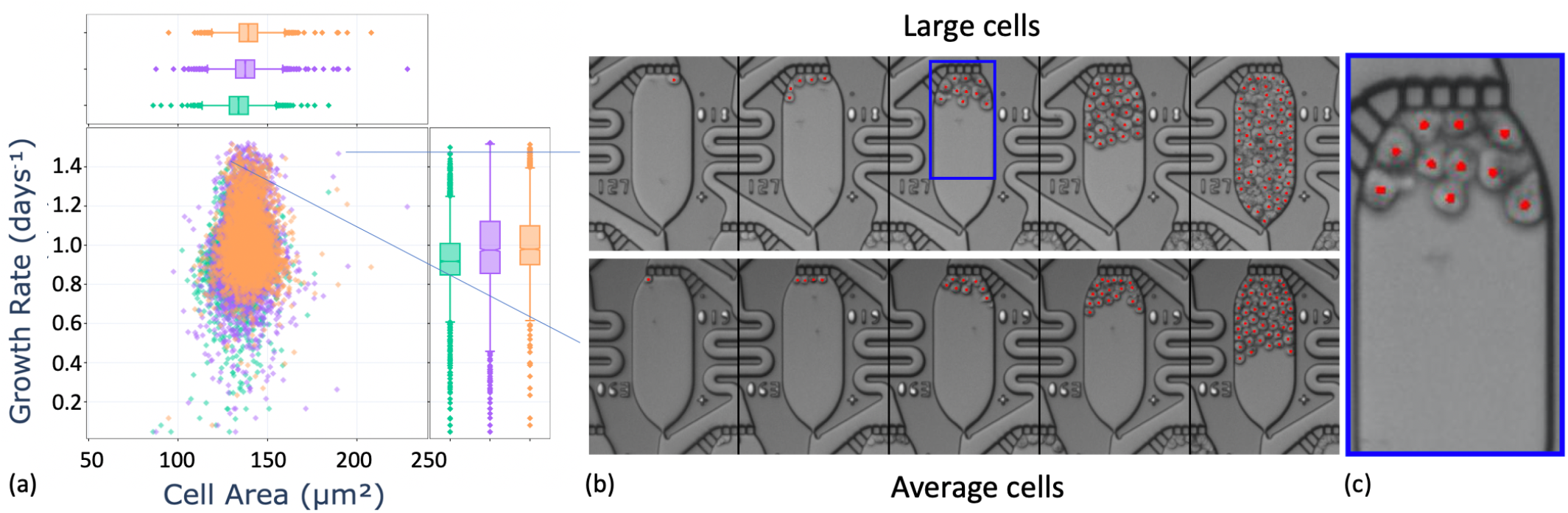
Phenotypic heterogeneity measured in extended duration culture. MOLM-13 cells were grown for 96 hr under continuous perfusion with RPMI media with 10% FBS. The growth rate distribution vs. time-averaged cell area across three chips (a), with example views of the average cell morphology compared to a rare subset of significantly larger cells (b), which were present at frequencies of approximately 0.05%. The time lapse images are taken at 24 hr intervals, and red dots are added to the images to depict the locations of the cell centroids as identified by the image analysis software. A higher magnification view of the cells with pear-shaped morphology is also shown (c).

### *In situ* fluorescence staining extends capabilities of high-throughput single-cell culture

In addition to time-resolved studies of clonal growth rates, the microfluidic platform is readily adapted for fluorescence imaging studies, including *in situ* live cell staining. In one demonstration, MOLM-13 cells were treated with a cell membrane-permeable nuclear stain (Hoescht 33258) as well as a PE-conjugated antibody against CD45, a marker expressed on all hematopoietic cells (Fig. 4a). Paired with Mask-RCNN cell instance segmentation, this experimental design can illuminate aspects of individual cell morphology and biomarker expression at high throughput on a clonal basis, including clonal phenotypic diversity. As an example, individual cell stain intensity distributions and signal statistics can be extracted for chambers of interest to study quantitative phenotypic differences within or across clones (Fig. 4b,c).

**Figure 4.**
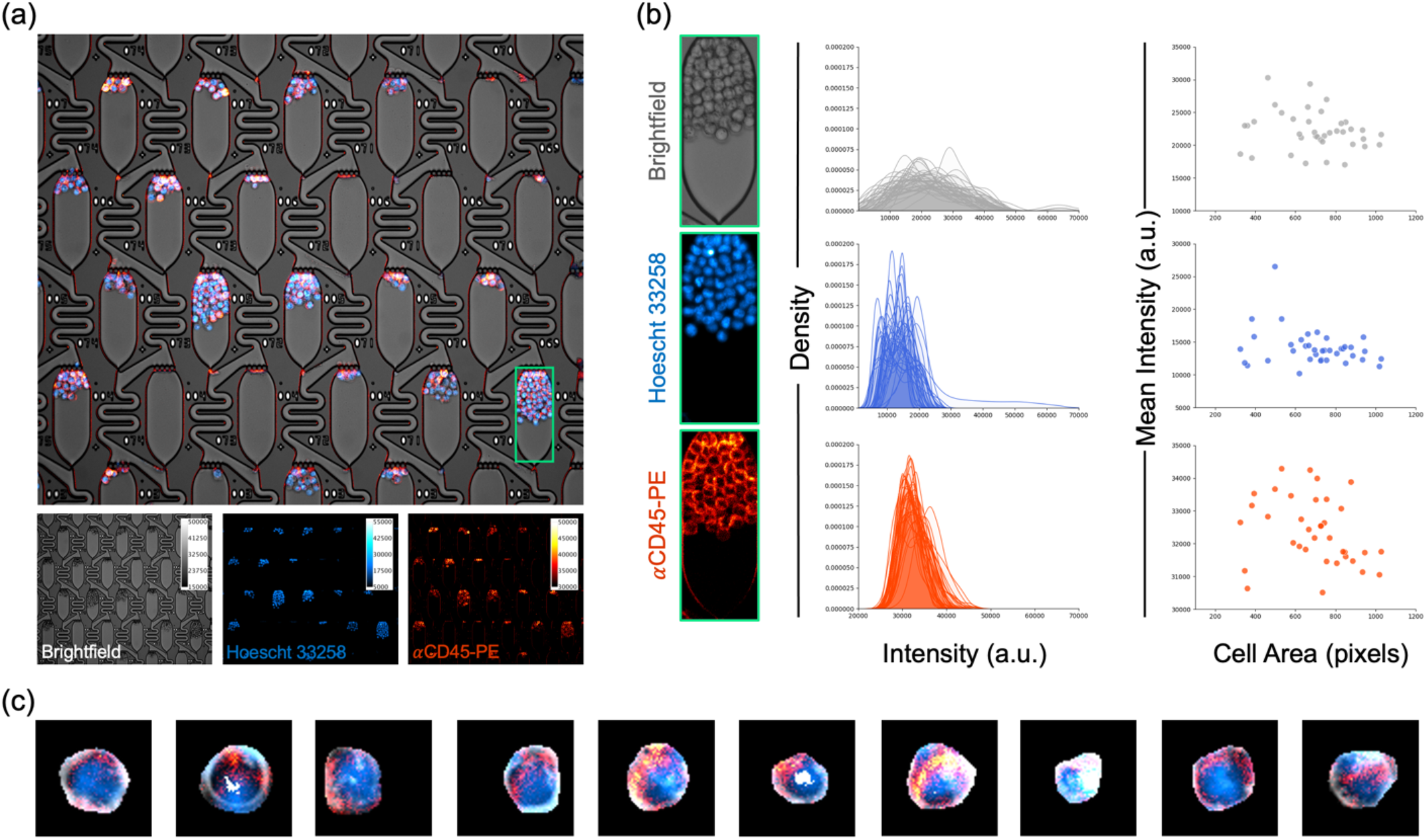
High-throughput extraction of clonal fluorescence data. (a) Multi-channel live-cell imaging of MOLM-13 clonal populations. After 72 hr of culture under constant flow, cells were stained *in situ* with Hoescht 33258 to visualize nuclei and PE-conjugated antibody against CD45, a pan-hematopoietic cell surface marker. (b) Signal quantification of MOLM-13 cells within a single culture chamber. Density plots depict brightfield, nuclear, and surface marker stain intensity distributions of automatically segmented individual cells within the selected chamber. Scatter plots present the relationship between cell size and mean intensity in each imaging channel. (c) Example multichannel images demonstrating diversity of individual cells segmented using Mask-RCNN.

### Rare drug resistant phenotypes are observed in multi-day cell culture experiments

The power of this platform to analyze thousands of cells per chip and many chips per system makes it uniquely suited for drug screening applications that require single cell resolution. To demonstrate this approach, we conducted an 8-chip study of cells exposed to either DMSO or 0.5 nM or 1.5 nM of the FLT3 inhibitor quizartinib (AC220). MOLM-13 cells harbor the internal tandem duplication (ITD) in-frame insertion in *FLT3*, a gene mutated in ∼30 percent of AML patients and associated with poor prognosis.[28] ITD renders FLT3 hyperactive via ligand independent phosphorylation, thus MOLM-13 cells are exquisitely sensitive to quizartinib.[29]

During long-term culture, the flow through the chip needs to be fast enough so that the metabolic waste products from upstream apartments do not significantly affect the downstream apartments. The vacuum pressure required to achieve an optimal flow rate was found to be in the range of -30 to -70mbar, depending on the total number of cells in the chip. When exposed to a vacuum pressure of -50mbar, the heatmaps in Fig. 5a reveal no apparent systematic bias in cell behavior across the chip, such as differing growth rates at positions nearer to the inlet versus the outlet. This finding supports the assumption that the growth properties of the single cells can be treated as statistically independent with regards to position inside the array. As expected, the cells thrived in the DMSO control, and a smaller fraction still grew well at the 0.5 nM conditions; however very few cells survived the 1.5 nM conditions. In the 0.5 nM conditions, the median growth rates per chip were reduced to 0.55 doublings/day with the middle growth rate quartiles falling in the range of 0.25 – 0.79 doublings/day. Surprisingly, we still observed fast-growing cells in the drugged condition that displayed growth rates up to 1.4 doublings/day – these rare cells appear to be practically unaffected by the drug treatment (Fig. 5b). One striking example of a drug resistant cell growing in the background of drug sensitive cells is shown in Fig. 5c after several days of exposure to 0.75 nM quizartinib (see Movie S1 and Fig. 5c).

**Figure 5.**
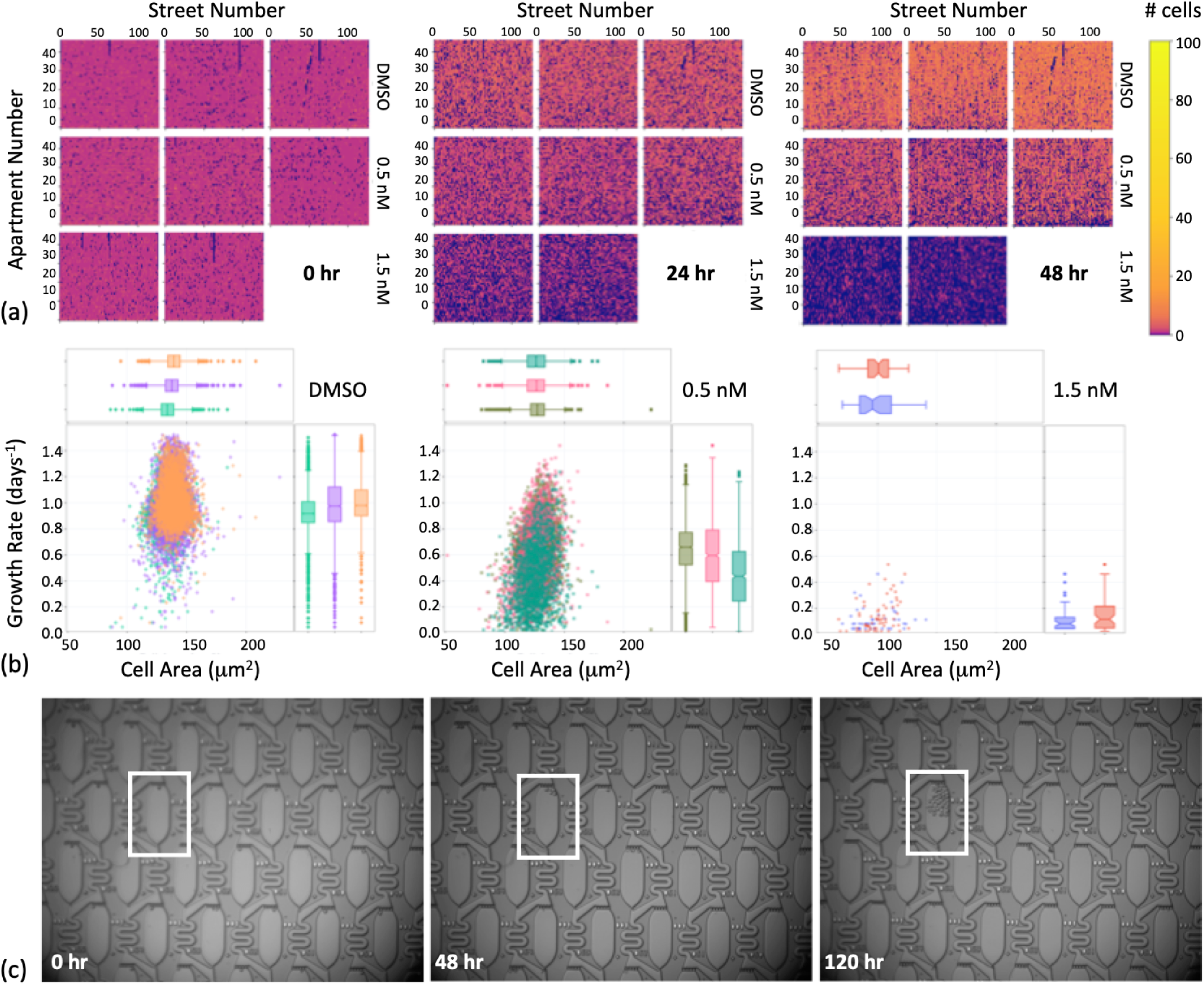
Growth rate heterogeneity due to drug response. The growth rates of MOLM-13 cells were measured in DMSO or 0.5 nM or 1.5 nM quizartinib over 96 hr. (a) To visualize the distribution of growth rates at different locations in the chip, the cell number per apartment is plotted a heatmap at t = 0, t = 24, and t = 48 hr. The heatmap colors are plotted on a log scale to better visualize the apartments with 0 or 1 cells. (b) The growth rates are shown in several scatter plots for the three cohorts depicting the relationship between doubling rate and mean cell area in each clone. (c) A time lapse of a single drug resistant clone emerging over 120 hr in 0.75 nM quizartinib is highlighted.

We also observed a consistent, positive correlation between cell area and growth rates across the different drug conditions, likely reflecting FLT-3 ITD’s established control over cell size and proliferation regulators like the mTOR and ERK pathways, respectively.[28] For example, the time-averaged median area per cell that was measured in the DMSO conditions was found to be 137 µm^2^ with the middle quartiles ranging from 128 – 144 µm^2^, whereas at 0.5 nM quizartinib the mean cell areas were reduced to a median of 126 µm^2^ and with the middle quartiles ranging from 119 – 133 µm^2^. However, we did not observe similar trends in the relationship between cell shape (eccentricity vs growth rate) as shown in Fig. S2. These relationships are further exemplified in Fig. S3, which shows a parallel coordinates plot linking the individual cell trajectories to the size dependence. The similarity of the growth trajectories across different cohorts was also classified with t-SNE plots in Fig. S4.

## Discussion

We have developed a high throughput live cell biology platform that can establish and maintain highly reproducible cellular architectures on chip for multiple days. This platform enables the analysis of phenotypic heterogeneity at the necessary scales for measuring low frequency variants in a population, such as cells that are resistant to a drug or have other rare morphological features. There are potential areas for improving this platform, such as by functionalizing the substrates with adhesive vs. non-adhesive patches at selective positions in the device and by using frit structures based on porous hydrogels,[30] which can help support and better constrain adherent and suspension cell cultures. We also expect that in the future these high-throughput phenotyping capabilities can be combined with the selective patterning of DNA primers inside the apartments to enable highly parallel transcriptome measurements to be perform in parallel with the image-based phenotyping for potential applications in single cell functional pharmacogenomics assays.

## Materials and Methods

### Experimental Design

#### Chip Fabrication

Microfluidic chips are fabricated on 6” wafers using deep reactive ion etching (DRIE) to form the channel walls, as previously described [31,32]. Photoresist (Shipley 1813) is spun onto the wafers at 500rpm for 5s and 4000 rpm for 60 s, baked at 115 °C for 60 s, exposed to 80-100 mJ/cm^2^ in a Karl Suss MA6 mask aligner, and then developed in Microposit MF319 developer for 30 s. The wafers are then thoroughly cleaned and etched to a depth of 15 – 20 µm in the DRIE (SPTS Pegasus Deep Silicon Etcher). The photoresist mask is then stripped and cleaned in piranha solution (3:1 H_2_SO_4_ to H_2_O_2_ at 200°C). Next, a 15µm thick layer of AZ 9260 photoresist is spun onto the backside of the wafer at 500 rpm for 5s and 1800rpm for 60s, baked at 110 °C for 60s, exposed to 4000 mJ/cm^2^ and developed for 300s in AZ400K 1:4 developer. This layer is used to create through silicon vias to establish the inlets and outlets and dice the chips. The photoresist is then stripped and thoroughly cleaned as described previously. Finally, we anodically bond borosilicate glass to the Silicon microchannels at 300 °C for 3 hr. In total, each wafer yields 12 devices (chips), which have dimensions of 30 mm X 25 mm.

#### Microfluidic Setup

Custom-made chip holders were machined in Aluminum (Protolabs, MN), comprising a bottom holder and a top-viewing window. The bottom piece contained ¼”-28 threaded holes to allow for connection to be made to the chips with screw-in luer locks (part number, company, city). The chip holders were also anodized (Surtronics, Raleigh, NC) to ensure that they would last inside the high humidity environment of a cell culture incubator for long durations. The chip holders were placed onto a custom stage adapter and mounted on a ASI-RAMM microscope (Applied Scientific Instrumentation, Eugene, OR), that contains an automated focus drive, objective changer, and filter changer. Fluid was introduced to the chip with an Elvesys pressure controller (OB1 MK3+, Paris, France) that applied vacuum pressure at the outlet.

#### Cell Culture

MOLM-13 acute myeloid leukemia cells [33] were obtained from the Wood lab. Cells were maintained in RPMI 1640 medium (Gibco 11875-093) supplemented with 10% fetal bovine serum (Gibco 10347-028) and penicillin/streptomycin (Gibco 15140-122) in a 5% CO_2_ environment. Cells were passaged in T25 flasks and centrifuged for 5 min at 350 rcf prior to sub-culturing to maintain a density range of 2.0 – 3.0 × 10^6^ cells per mL. A new thaw of cells was utilized every 8 weeks to minimize genetic drift. Counting and viability determination with 0.4% Trypan Blue with a Countess II instrument (ThermoFisher Scientific). Quizartinib (AC220) was obtained from Selleck Chemicals LLC.

#### Cell Loading

Cells were loaded into the chip by pipetting a 20mL aliquot into screw-in luer locks positioned on the inlet side, after which the cells were infused into the chip by applying 20 30 mbar vacuum pressure to the outlet side using a syringe body that was attached to a rubber stopper. The microfluidic architecture consists of one inlet and one outlet, which feed into the active area of the chip by successive flow division in a binary tree, leading to 128 parallel streets with 47 apartments in series. The loading time typically required 3 – 5 minutes for the cells to reach the last row of apartments in each street, corresponding to a loading rate of about 20 cells per second. After the cells were trapped in each constriction, the luer locks on the inlet side were rinsed at least 3 times by replacing the fluid with fresh cell culture media. In order to eliminate any remaining cells that were stuck in the luer lock or on the chip surface, we irradiated the luer locks with UVC using a 270nm LED diode attached to a heat sink (Irtronix, Torrance, CA) – this provided a lethal radiation dose to any non-specifically adhered cells and prevented the chips from being invaded with cells at later time points. Finally, the cells were squeezed through the constrictions by applying a brief (∼1s) pressure pulse in the range of 300 − 800mbar to the outlet. The chips were then disconnected from the imager and put into the incubator.

#### High-Throughput Microscopy

We developed custom python codes to rapidly take images of each chamber. The algorithm involved first identifying 3 crosshairs on the chip to establish the equation of a plane, next creating a stage position list containing the XY position and optimal focal plane for each image, then taking images of each chamber, and finally saving and naming the images in custom formats to render them compatible with the computer vision algorithms. The software used to image the chips is provided at github,[26] and since they are based on a python wrapper for Micro-Manager [34], the program is easily adapted for most standard robotic microscopes.

#### Fluorescence Imaging

Chips were loaded with MOLM-13 cells and cultured in 0.2 µm-filtered R10 media at 37°C and 5% CO_2_. At 72 hr, 5 µL of Hoescht 33258 (0.1 mg/mL) was added to ∼50 µL of media at the microfluidic inlet port and flowed onto the chip using negative pressure (-100 mbar) applied at the microfluidic outlet. Constant flow was maintained for 10-15 minutes to stain cell nuclei, followed by rinsing with media. Similarly, 5 µL of PE-conjugated anti-CD45 (Invitrogen, 0.2 mg/mL) was flowed in, incubated, and rinsed prior to imaging. Multichannel images were collected in brightfield and using standard DAPI and Texas Red filter sets.

### Statistical Analysis

#### Image Analysis

We developed custom python codes to rapidly analyze the images and extract cellular phenotypic properties in a computationally efficient manner. Our cell extraction algorithms make use of the Mask-RCNN image segmentation model, [35] which is designed to identify objects in images without the need for pixel classification post-processing. This is an advance on previous methods for biological image segmentation [36,37] that enabled us to compose a simple pipeline for cell quantification using a minimal amount of training data. In a separate study, we quantified the superior performance of Mask-RCNN segmentation relative to supervised segmentation algorithms and statistical methods.[38] Similarly, the SVHN (street view housing number)[39] model is an architecture for digit classification that we used to determine the apartment identifiers etched into the chips.

Our pipeline consists of three separately applied models where the first is used to identify a key point within each apartment image (hereto referred to as a “marker”), given an image containing multiple apartments (i.e. raw microscope images). Images of individual apartments were then extracted using these markers. Because the raw microscope images often have slight rotations, the relative positions of the identified markers in adjacent apartments were used to infer an overall rotation of the images to be inverted before further decomposing the individual apartment images. The apartment images were then registered against a template image to remove small translations. The digit identifiers for each apartment, with no rotations or translations, were extracted based on fixed offsets from the marker position. Fixed offsets are determined relative to several chip landmarks and need to be updated whenever the chip form factor is altered. Identification of individual cell objects is performed based on the entire apartment image, but segmented results are filtered to the chamber and trap areas, again using fixed offsets from the marker, as a way to prohibit erroneous classification of debris within microfluidic channels.

Training for the cell segmentation model included 814 annotated images and the Mask-RCNN model trained was initialized to a weight set resulting from pre-training over the COCO [40] image dataset, a feature provided by the Matterport implementation.[25] Training also included an augmentation pipeline consisting of image flips, affine rotations, random croppings, contrast transform, and blurring. The marker identification model was trained in a very similar fashion but required only 70 annotated images since the associated classification task was simpler. By contrast, the digit recognition model required far more training data (9,375 annotated images) though this annotation task was much less time consuming since the individual digit images only needed to be assigned a class, as such bounding boxes or object masks were not required.

We have also developed a dashboard visualization tool that allows the growth rates and other properties to be viewed at the experiment level, individual apartment level, and array levels. More details on the software package can be found at github.[22]

#### Data Analysis

The data presented in Figures 3 & 5 is limited to chambers starting from a single cell and having at least one cell in the apartment at each time point. This led to significantly fewer data points for the 1.5 nM quizartinib cohort, where a majority of cells did not survive the drug treatment over several days. The growth rates are determined by fitting the raw trajectories to an exponential with base 2. The calculated growth rates are likely to be a lower bound, since the image-segmentation models begin to miss cells in apartments that are very crowded, as shown in Fig. 3b.

## H2: Supplementary Materials

Supplementary Theory

Fig. S1. Experimental setup

Fig. S2. Scatterplot of growth rate versus cell size or shape

Fig. S3. Parallel coordinates plot of growth trajectories and mean cell areas

Fig. S4. t-SNE plot of growth trajectories

Movie S1. Drug resistant AML clone growing in 0.75 nM quizartinib

## General

The authors are thankful to prior members of the Yellen lab who contributed to early experiments and testing of microfluidic designs.

## Author contributions

B.B.Y. designed and fabricated the chips, built the imaging and pumping systems, wrote early versions of the automated imaging codes, trained the cell classifier, and analyzed the datasets. J.S.Z. performed and analyzed the drug resistance experiments. E.A.C. wrote the data visualization software package, trained the digit and marker classifiers, and maintains the software on a github repository. C.I.S. wrote the fully automated imaging codes and maintains the software on a github repository. E.D.S. conducted the biological experiments with *in situ* fluorescent imaging. Z.G.F., M.A.L., K.C.W., and J.H. supervised the experiments and data analysis methods. All authors provided comments to the manuscript. All authors have seen and approved the manuscript, which has not been accepted or published elsewhere.

## Data and materials availability

All data needed to evaluate the conclusions in the paper are present in the paper and/or the Supplementary Materials.

## Supplementary Theory

Laminar flow hydrodynamic networks can be modeled like electrical circuits, where the pressure, flow rate, and hydrodynamic resistances are analogous to voltage, current, and electrical resistances. Ladder (or mesh) networks are comprised of two types of resistors, including those aligned parallel to the main flow path, i.e., R_A_ and R_T_, (the rails of the ladder) and those aligned perpendicular to the main flow path, i.e., R_B_ (the rungs of the ladder). The flow distribution can be solved by setting up continuity equations at each branch point in the array. From there, we apply a constant pressure drop, ΔP, parallel to the flow direction across each array period. This system of equations thus reduces to solving the pressure at 4 nodes in the minimum unit cell, which are given by:

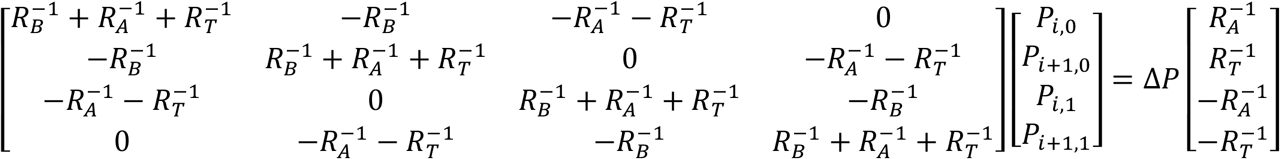

where P_i, 0_, P_i,1_, P_i+1,0_, and P_i+1,1_ are the four unique nodes of the unit lattice.

The pressures at each node can be solved by inverting Eq. (1) to yield a generic solution in terms of the pressure at an arbitrary point, in this case chosen as P_i,0_:

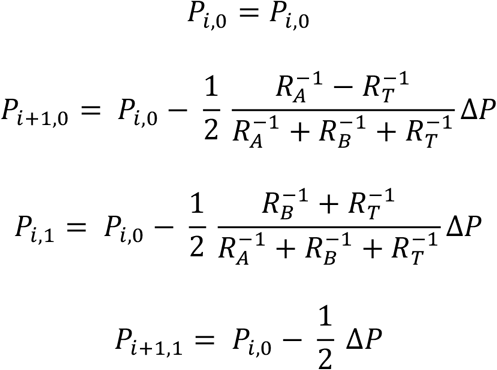

We can then determine the ratio of flow along the two lateral paths, *Q*_*B*_, relative to the flow through the apartment, *Q*_*A*_, which are given by:

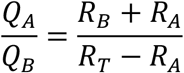

Since this ratio changes sign as a function of the relative magnitude of R_A_ and R_T_, this result indicates that there are two regimes of fluid flow. When R_T_ > R_A_, which is the typical scenario for previously studied trapping designs, the flow ratio is positive and approaches a singularity when R_T_ is nearly equal to R_A_. This singularity defines a critical point where there is zero flow through the lateral branches, R_B_, and all of the flow moves solely through the R_A_ and R_T_ paths, practically in straight lines. An alternative way to think of this phenomenon is that the pressure at the adjacent nodes P_i,0_ and P_i,1_ are equal when R_A_ and R_T_ have equal resistance, leading to zero flow in the lateral branches.

The other flow regime, which has not previously been reported, occurs when R_T_ < R_A_, which leads to the ratio in Eq. (3) becoming negative. The significance of this sign inversion is that the flow through the lateral branches, *Q*_*B*_, is assigned in the wrong direction. In this flow regime, all of the fluid joins together at the branch point and flows through the trap, which is a perfect trap from a mathematical sense.

## Supplementary Figures

**Figure S1.**
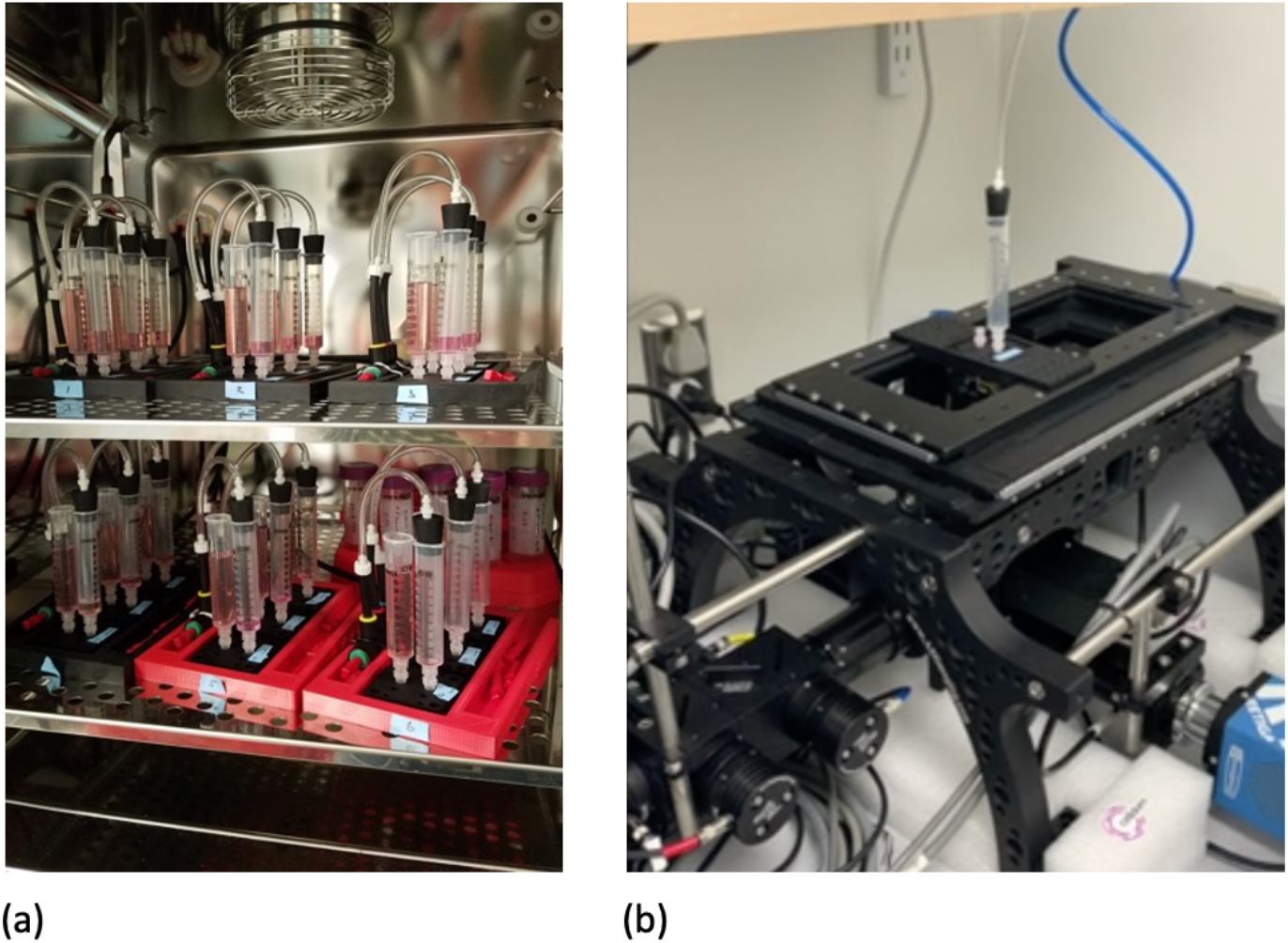
The incubator system used for maintaining chips with media is shown (a). Continuous media flow through the chips was achieved by pulling vacuum at the outlet, while allowing the media at the inlet to be exposed to the gas- and temperature-controlled environment of the incubator. External to the environment, the pressure was maintained by 6 vacuum regulators, which were divided with 3-way splitter manifolds, enabling up to 18 chips to be contained simultaneously. The chip on the imager is shown (b).

**Figure S2.**
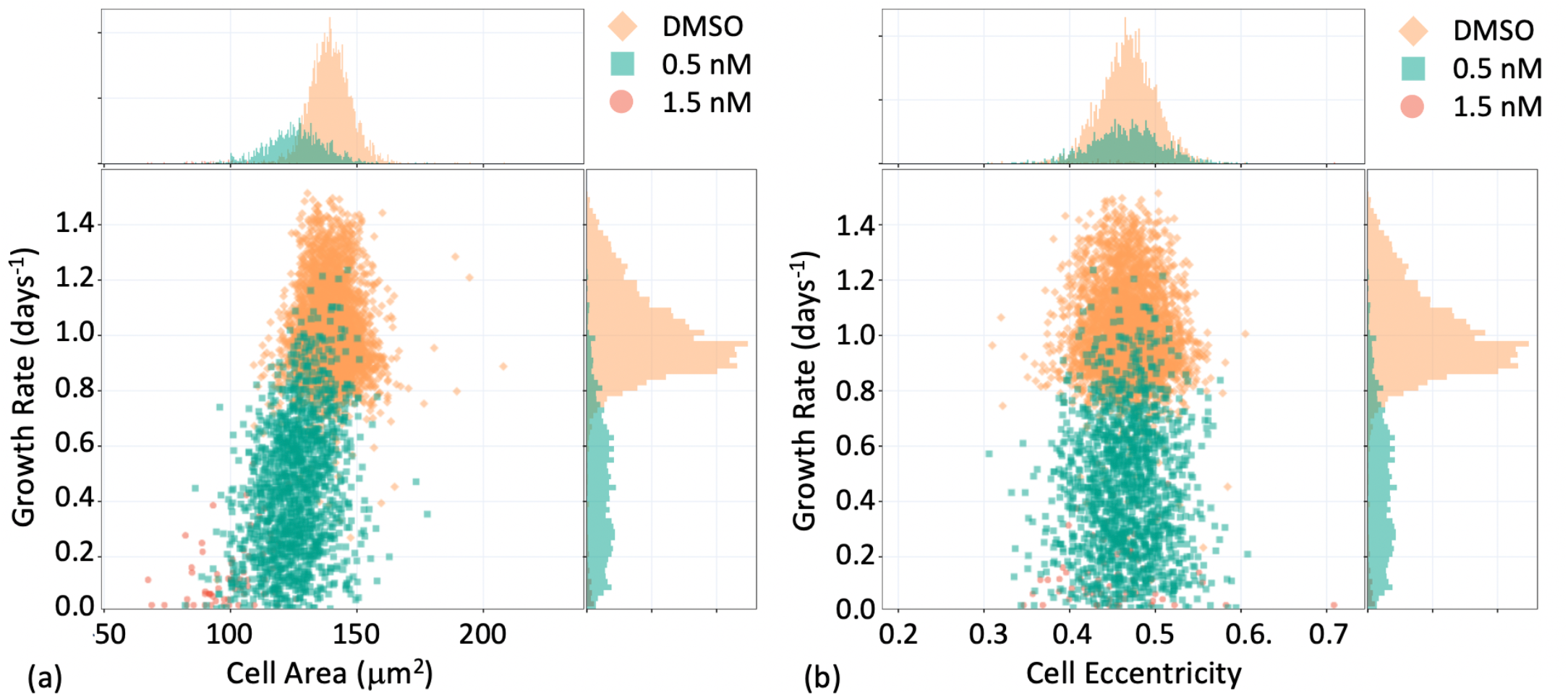
Size and shape dependence on clonal growth rates is shown for an example chip in the 1.5 nM, 0.5 nM and DMSO cohort. There is positive correlation between cell size and growth rate, however a correlation between cell shape and growth rate was not observed.

**Figure S3.**
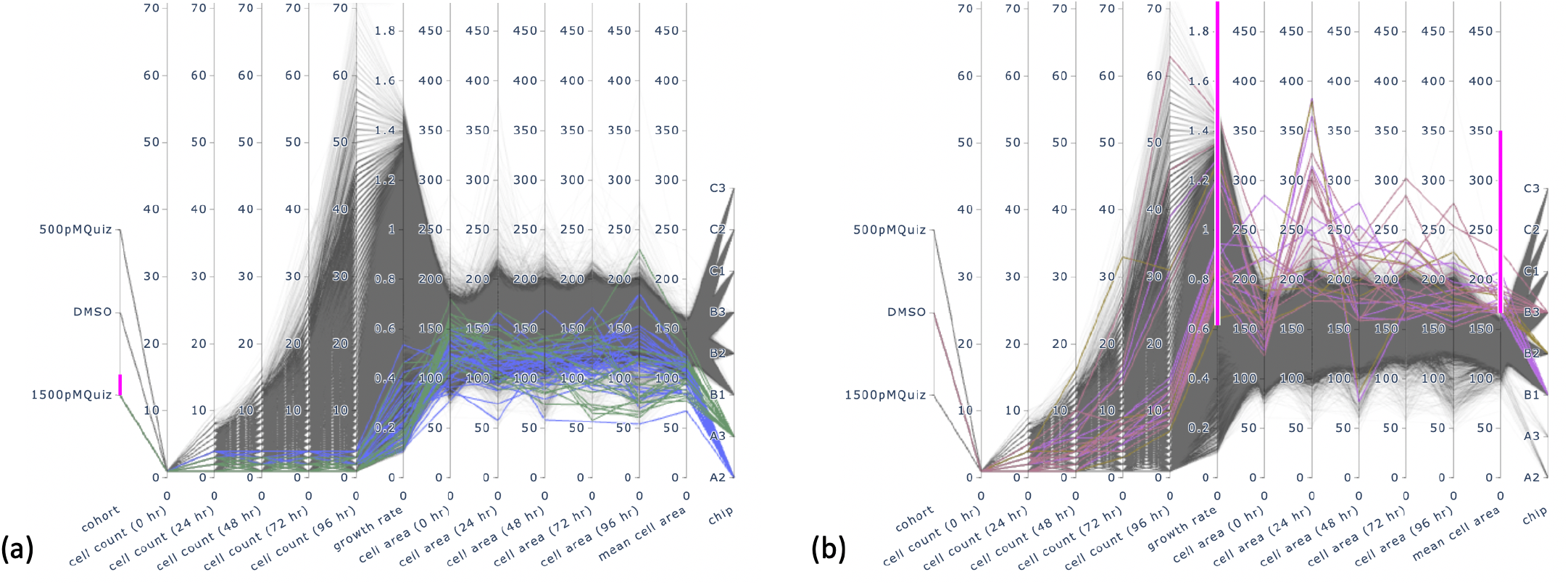
The parallel coordinates plot shows the growth and size trajectory for each cell identified in the different cohorts and chips and is used as compact method for visualizing linked phenotypic attributes. The pink bars represent the filtering to highlight the linked attributes. (a) The growth trajectories of the 1.5 nM quizartinib cohort, which are highlighted in the blue and green trajectories, show significantly lower growth rates and smaller time-averaged cell size. (b) A subset of cells are filtered to display the cell trajectories with growth rates greater than 0.66 doublings per day and with cell areas greater than 150 µm^2^.

**Figure S4.**
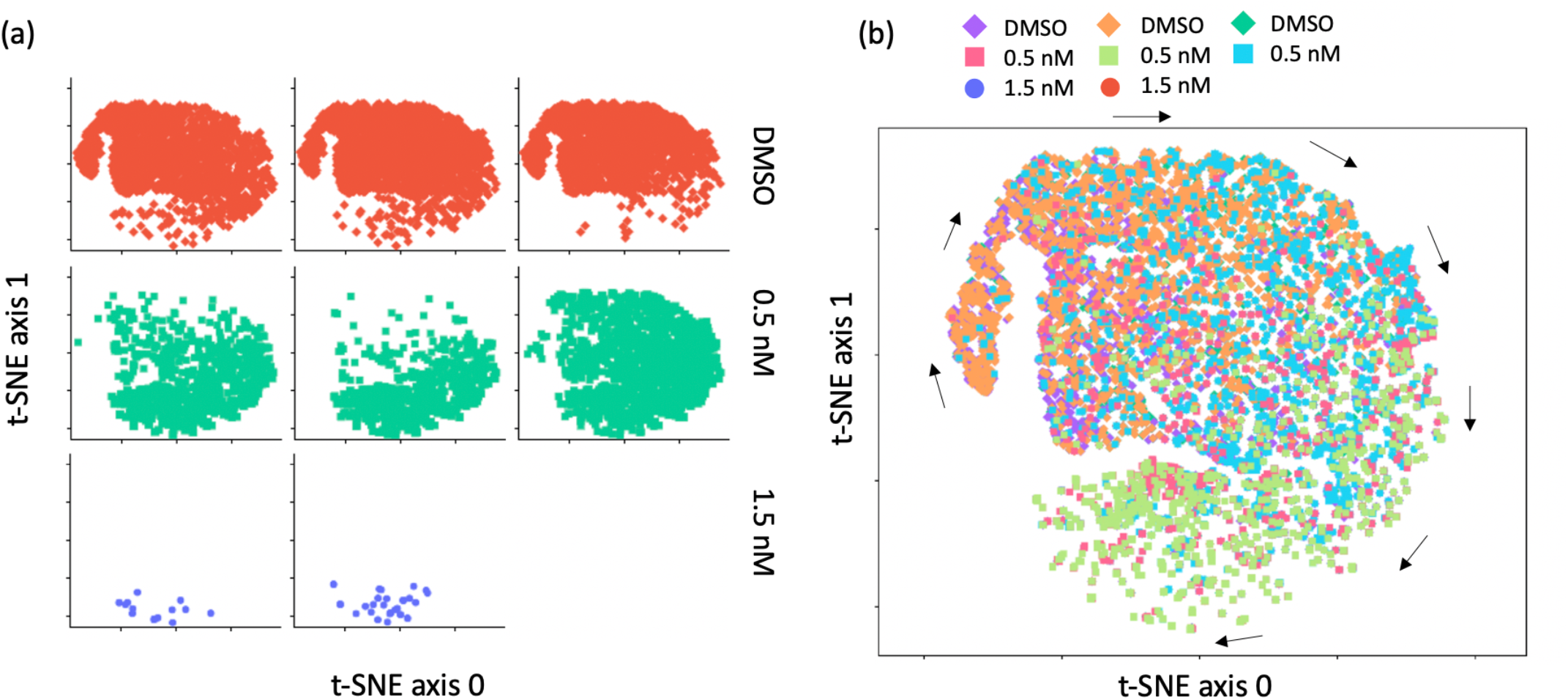
t-SNE plot is shown for the cell trajectory data. The arrows denote the direction of decreasing cell doubling rate. The individual cluster map for each chip is shown in (a) and the combined data overlaid is shown in (b).

## Supplementary Movies

**Movie S1.**
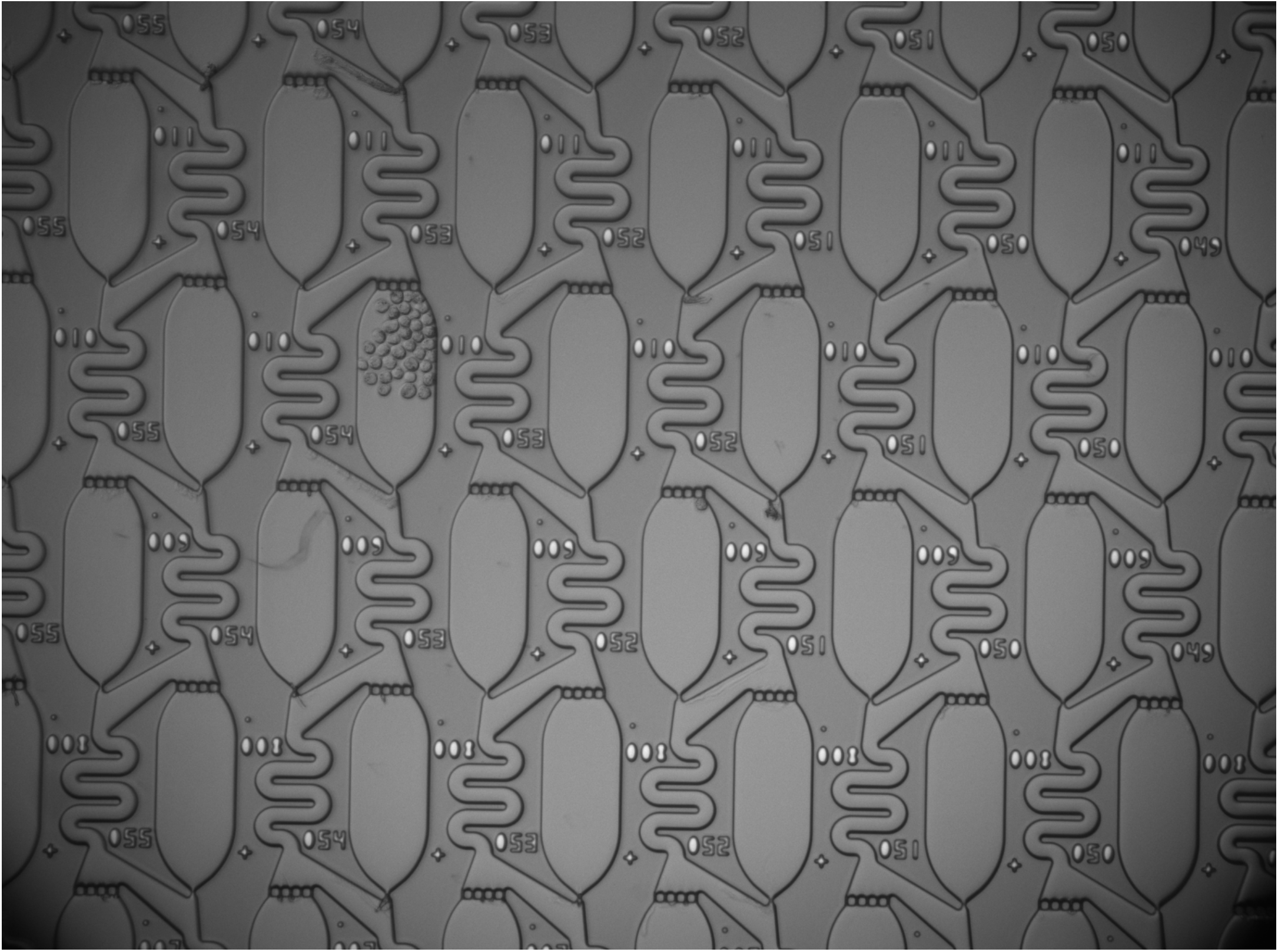
This time-lapse video shows a single drug-resistant MOLM-13 clone emerging after 120 hr of continuous exposure to 0.75 nM quizartinib.

## References

1. Kim C, Gao R, Sei E, Brandt R, Hartman J, Hatschek T, Crosetto N, Foukakis T, Navin NE (2018) Chemoresistance Evolution in Triple-Negative Breast Cancer Delineated by Single-Cell Sequencing. Cell 173: 879-893.e813.

2. Evrony Gilad D, Lee E, Mehta Bhaven K, Benjamini Y, Johnson Robert M, Cai X, Yang L, Haseley P, Lehmann Hillel S, Park Peter J, Walsh Christopher A (2015) Cell Lineage Analysis in Human Brain Using Endogenous Retroelements. Neuron 85: 49–59.

3. Tanay A, Regev A (2017) Scaling single-cell genomics from phenomenology to mechanism. Nature 541: 331–338.

4. Regev A, Teichmann SA, Lander ES, Amit I, Benoist C, Birney E, Bodenmiller B, Campbell P, Carninci P, Clatworthy M, Clevers H, Deplancke B, Dunham I, Eberwine J, Eils R, Enard W, Farmer A, Fugger L, Göttgens B, Hacohen N, Haniffa M, Hemberg M, Kim S, Klenerman P, Kriegstein A, Lein E, Linnarsson S, Lundberg E, Lundeberg J, Majumder P, Marioni JC, Merad M, Mhlanga M, Nawijn M, Netea M, Nolan G, Pe’er D, Phillipakis A, Ponting CP, Quake S, Reik W, Rozenblatt-Rosen O, Sanes J, Satija R, Schumacher TN, Shalek A, Shapiro E, Sharma P, Shin JW, Stegle O, Stratton M, Stubbington MJT, Theis FJ, Uhlen M, van Oudenaarden A, Wagner A, Watt F, Weissman J, Wold B, Xavier R, Yosef N, Human Cell Atlas Meeting P (2017) The Human Cell Atlas. eLife 6: e27041.

5. Petti AA, Williams SR, Miller CA, Fiddes IT, Srivatsan SN, Chen DY, Fronick CC, Fulton RS, Church DM, Ley TJ (2019) A general approach for detecting expressed mutations in AML cells using single cell RNA-sequencing. Nature Communications 10: 3660.

6. Zheng GXY, Terry JM, Belgrader P, Ryvkin P, Bent ZW, Wilson R, Ziraldo SB, Wheeler TD, McDermott GP, Zhu J, Gregory MT, Shuga J, Montesclaros L, Underwood JG, Masquelier DA, Nishimura SY, Schnall-Levin M, Wyatt PW, Hindson CM, Bharadwaj R, Wong A, Ness KD, Beppu LW, Deeg HJ, McFarland C, Loeb KR, Valente WJ, Ericson NG, Stevens EA, Radich JP, Mikkelsen TS, Hindson BJ, Bielas JH (2017) Massively parallel digital transcriptional profiling of single cells. 8: 14049.

7. Stuart T, Satija R (2019) Integrative single-cell analysis. Nature Reviews Genetics 20: 257–272.

8. Beaumont KG, Hamou W, Bozinovic N, Silvers TR, Shah H, Dave A, Allette K, Strahl M, Wang Y-c, Arib H, Antoine A, Ellis E, Smith M, Bruhn B, Dottino P, Martignetti JA, Schadt E, White M, Sebra R (2018) Multiparameter cell characterization using nanofluidic technology facilitates real-time phenotypic and genotypic elucidation of intratumor heterogeneity. bioRxiv: 457010.

9. Mocciaro A, Roth TL, Bennett HM, Soumillon M, Shah A, Hiatt J, Chapman K, Marson A, Lavieu G (2018) Light-activated cell identification and sorting (LACIS) for selection of edited clones on a nanofluidic device. Communications Biology 1: 41.

10. Amirouchene-Angelozzi N, Swanton C, Bardelli A (2017) Tumor Evolution as a Therapeutic Target. Cancer Discovery.

11. Chatterjee N, Bivona TG (2019) Polytherapy and Targeted Cancer Drug Resistance. Trends in Cancer 5: 170–182.

12. McCoach CE, Bivona TG (2019) Engineering Multidimensional Evolutionary Forces to Combat Cancer. Cancer Discovery 9: 587.

13. Skelley AM, Kirak O, Suh HY, Jaenisch R, Voldman J (2009) Microfludic control of cell pairing and fusion. Nature Methods 6: 147–152.

14. Di Carlo D, Wu LY, Lee LP (2006) Dynamic single cell culture array. Lab on a Chip 6: 1445–1449.

15. Jin D, Deng B, Li JX, Cai W, Tu L, Chen J, Wu Q, Wang WH (2015) A microfluidic device enabling high-efficiency single cell trapping. Biomicrofluidics 9: 014101.

16. Crane MM, Clark IBN, Bakker E, Smith S, Swain PS (2014) A Microfluidic System for Studying Ageing and Dynamic Single-Cell Responses in Budding Yeast. PLOS ONE 9: e100042.

17. Dura B, Dougan SK, Barisa M, Hoehl MM, Lo CT, Ploegh HL, Voldman J (2015) Profiling lymphocyte interactions at the single-cell level by microfluidic cell pairing. Nature Communications 6: 5940.

18. Islam M, Brink H, Blanche S, DiPrete C, Bongiorno T, Stone N, Liu A, Philip A, Wang G, Lam W, Alexeev A, Waller EK, Sulchek T (2017) Microfluidic Sorting of Cells by Viability Based on Differences in Cell Stiffness. Scientific Reports 7: 1997.

19. DiTommaso T, Cole JM, Cassereau L, Buggé JA, Hanson JLS, Bridgen DT, Stokes BD, Loughhead SM, Beutel BA, Gilbert JB, Nussbaum K, Sorrentino A, Toggweiler J, Schmidt T, Gyuelveszi G, Bernstein H, Sharei A (2018) Cell engineering with microfluidic squeezing preserves functionality of primary immune cells in vivo. Proceedings of the National Academy of Sciences 115: E10907.

20. Sharei A, Zoldan J, Adamo A, Sim WY, Cho N, Jackson E, Mao S, Schneider S, Han M-J, Lytton-Jean A, Basto PA, Jhunjhunwala S, Lee J, Heller DA, Kang JW, Hartoularos GC, Kim K-S, Anderson DG, Langer R, Jensen KF (2013) A vector-free microfluidic platform for intracellular delivery. Proceedings of the National Academy of Sciences 110: 2082.

21. Caicedo JC, Goodman A, Karhohs KW, Cimini BA, Ackerman J, Haghighi M, Heng C, Becker T, Doan M, McQuin C, Rohban M, Singh S, Carpenter AE (2019) Nucleus segmentation across imaging experiments: the 2018 Data Science Bowl. Nature Methods 16: 1247–1253.

22. Czech EA (2020) hammerlab/SmartCount: v1.0.0. doi:105281/zenodo4304993.

23. Li Y, Jang JH, Wang C, He B, Zhang K, Zhang P, Vu T, Qin L (2017) Microfluidics Cell Loading-Dock System: Ordered Cellular Array for Dynamic Lymphocyte-Communication Study. Advanced Biosystems 1: 1700085.

24. Yang SJ, Berndl M, Michael Ando D, Barch M, Narayanaswamy A, Christiansen E, Hoyer S, Roat C, Hung J, Rueden CT, Shankar A, Finkbeiner S, Nelson P (2018) Assessing microscope image focus quality with deep learning. BMC Bioinformatics 19: 77.

25. Abdulla W (2017) Mask R-CNN for object detection and instance segmentation on Keras and TensorFlow. Github repository: https://github.com/matterport/Mask_RCNN.

26. Sanford CI, Yellen BB (2020) yellenlab/SmartScope: v1.0. doi:105281/zenodo4319319.

27. Lloyd Alison C (2013) The Regulation of Cell Size. Cell 154: 1194–1205.

28. Daver N, Schlenk RF, Russell NH, Levis MJ (2019) Targeting FLT3 mutations in AML: review of current knowledge and evidence. Leukemia 33: 299–312.

29. Zarrinkar PP, Gunawardane RN, Cramer MD, Gardner MF, Brigham D, Belli B, Karaman MW, Pratz KW, Pallares G, Chao Q, Sprankle KG, Patel HK, Levis M, Armstrong RC, James J, Bhagwat SS (2009) AC220 is a uniquely potent and selective inhibitor of FLT3 for the treatment of acute myeloid leukemia (AML). Blood 114: 2984–2992.

30. Decock J, Schlenk M, Salmon J-B (2018) In situ photo-patterning of pressure-resistant hydrogel membranes with controlled permeabilities in PEGDA microfluidic channels. Lab on a Chip 18: 1075–1083.

31. Ohiri KA, Kelly ST, Motschman JD, Lin KH, Wood KC, Yellen BB (2018) An acoustofluidic trap and transfer approach for organizing a high density single cell array. Lab on a Chip 18: 2124–2133.

32. Li Y, Motschman JD, Kelly ST, Yellen BB (2020) Injection Molded Microfluidics for Establishing High-Density Single Cell Arrays in an Open Hydrogel Format. Analytical Chemistry 92: 2794–2801.

33. Matsuo Y, MacLeod RAF, Uphoff CC, Drexler HG, Nishizaki C, Katayama Y, Kimura G, Fujii N, Omoto E, Harada M, Orita K (1997) Two acute monocytic leukemia (AML-M5a) cell lines (MOLM-13 and MOLM-14) with interclonal phenotypic heterogeneity showing MLL-AF9 fusion resulting from an occult chromosome insertion, ins(11;9)(q23;p22p23). Leukemia 11: 1469–1477.

34. Edelstein A, Amodaj N, Hoover K, Vale R, Stuurman N (2010) Computer Control of Microscopes Using µManager. Current Protocols in Molecular Biology 92: 14.20.11-14.20.17.

35. He K, Gkioxari G, Dollar P, Girshick R (2017) Mask R-CNN. The IEEE International Conference on Computer Vision (ICCV). pp. 2961–2969.

36. Ronneberger O, Fischer P, Brox T (2015) U-Net: Convolutional Networks for Biomedical Image Segmentation. arXiv:150504597.

37. Berg S, Kutra D, Kroeger T, Straehle CN, Kausler BX, Haubold C, Schiegg M, Ales J, Beier T, Rudy M, Eren K, Cervantes JI, Xu B, Beuttenmueller F, Wolny A, Zhang C, Koethe U, Hamprecht FA, Kreshuk A (2019) ilastik: interactive machine learning for (bio)image analysis. Nature Methods 16: 1226–1232.

38. SoRelle ED WS, Yellen BB, Wood KC, Luftig MA, Chan C (2020) Comparing Instance Segmentation Methods for Analyzing Clonal Growth of Single Cells in Microfluidic Chips. bioRxiv: TBD.

39. Goodfellow IJ, Bulatov Y, Ibarz J, Arnoud S, Shet V (2013) Multi-digit Number Recognition from Street View Imagery using Deep Convolutional Neural Networks. arXiv:13126082.

40. Lin T-Y, Maire M, Belongie S, Bourdev L, Girshick R, Hays J, Perona P, Ramanan D, Zitnick CL, Dollar P (2014) Microsoft COCO: Common Objects in Context. arXiv:14050312

